# Transcription-driven establishment of DNA methylation at the PWS/AS imprinted region

**DOI:** 10.64898/2026.09.08.749716

**Authors:** Josephine Haake, Alexander Lederhof, Claudia Pommerenke, Laura Steenpass

## Abstract

Genomic imprinting is regulated by allele-specific DNA methylation at imprinting control regions, ensuring parent-of-origin-specific gene expression. The human Prader-Willi/Angelman syndrome (PWS/AS) locus on chromosome 15 harbours a bipartite imprinting center consisting of the PWS-SRO, which acquires maternal-specific methylation, and the AS-SRO, an upstream promoter that initiates transcription in oocytes. To directly test whether transcription across the PWS-SRO is sufficient to trigger de novo DNA methylation, a human iPSC based in vitro system was established in which the endogenous AS-SRO was replaced by a doxycycline-inducible promoter driving transcription across the PWS-SRO. Inducible transcription was robustly activated upon doxycycline treatment and confirmed by splicing to *SNRPN* exon 2, mirroring the structure of oocyte-derived transcripts. Gain of DNA methylation up to 10% was observed only following directed differentiation into the endodermal lineage. This study provides the first direct evidence in a human model that transcription across the PWS-SRO is necessary but not sufficient for de novo DNA methylation. Instead, differentiation-dependent cues, potentially resembling aspects of the oocyte environment, are required. The established in vitro system enables controlled dissection of the molecular factors and chromatin dynamics underlying imprint establishment at the PWS/AS locus.

**GRAPHICAL ABSTRACT:** 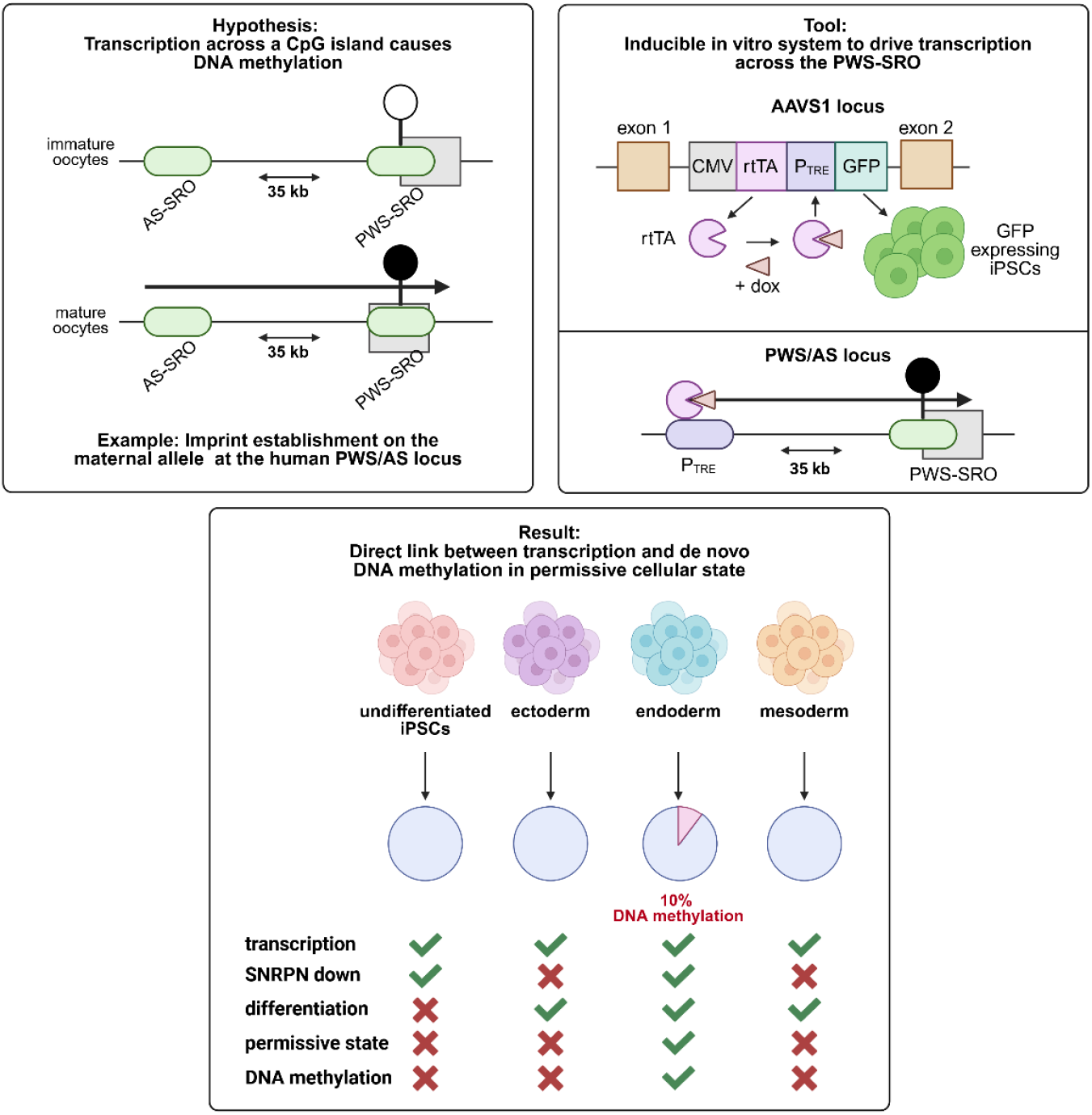

## INTRODUCTION

Maternal and paternal genomes are not functionally equivalent in mammals due to genomic imprinting, an epigenetic process where genes are silenced depending on their parental origin (1). Genomic imprinting is regulated by DNA methylation, whereby methyl groups are added to cytosine bases within CpG dinucleotides, typically leading to stable gene repression (2). DNA methylation is targeted to CpG-rich imprinting control regions (ICRs), which regulate parent-of-origin-specific gene expression through allele-specific methylation. One parental allele is methylated and transcriptionally silent, whereas the other remains unmethylated and active, yielding an average methylation level of ∼50%. (3). This methylation asymmetry defines the ICR as a germline differentially methylated region (gDMR). The regulation of genomic imprinting at gDMRs follows a defined life cycle consisting of three major phases: establishment, maintenance and erasure. During establishment, parent-specific DNA methylation patterns are set in the germline, specifically in oocytes and sperm at ICRs, ensuring that maternal and paternal alleles carry distinct epigenetic signatures (4–7). These methylation marks are then maintained throughout embryonic development and in somatic cells, despite the genome-wide demethylation events that occur after fertilization. This maintenance is essential for preserving monoallelic expression of imprinted genes (8). In the erasure phase, occurring during the development of primordial germ cells (PGCs) in the embryo, existing imprints are removed to allow for the establishment of new, sex-specific imprints in the next generation. This cyclical reprogramming ensures the proper transmission and regulation of imprinted genes across generations (9,10).

The imprinted Prader-Willi/Angelman syndrome locus (PWS/AS) on human chromosome 15q11-q13 has a bipartite imprinting center consisting of two functional elements: the PWS-SRO, which becomes methylated on the maternal allele during oocyte maturation, and the AS-SRO, located 35kb upstream of the PWS-SRO (SRO: shortest region of overlap; Supplementary Fig. S1A) (11). The AS-SRO is essential for DNA methylation establishment at the PWS-SRO, serving as an oocyte-specific promoter (12–14). Within the PWS/AS locus, most imprinted genes are expressed from the paternal allele, whereas *UBE3A* is predominantly expressed from the maternal allele in neurons (15). The PWS-SRO functions as the promoter of the ubiquitously expressed *SNURF-SNRPN* transcript and, through DNA methylation, regulates paternal expression of the upstream genes *MKRN3*, *MAGEL2*, *NDN* and *NPAP1* as well as the downstream snoRNA cluster *SNORD116* by a cis-acting mechanism that remains incompletely understood. Monoallelic expression of *MKRN3*, *MAGEL2* and *NDN* is further reinforced by maternally methylated secondary DMRs (16–19). In neurons, the paternally expressed long non-coding RNA *SNHG14*, extends across the *UBE3A* locus. Convergent transcription of *SNHG14* and *UBE3A* results in transcriptional interference, with RNA polymerase collisions in intron 4 resulting in downregulation of the paternal *UBE3A* allele (20,21). Loss of *UBE3A* expression from the maternal allele cannot be compensated in neurons and leads to Angelman syndrome (12,22), a neurodevelopmental disorder characterized by severe intellectual and developmental disabilities (23). The AS-SRO functions as a promoter during oocyte development, producing non-coding transcripts that traverse the PWS-SRO, a process thought to be critical for triggering de novo methylation on the maternal allele (13,24,25). There is increasing evidence that transcription plays a critical role in the establishment of genomic imprints. At the murine *Gnas* locus, transcription from the upstream *Nesp* promoter is required for de novo methylation of downstream maternal DMRs during oocyte development (26). Similar findings were made at the murine *Zrsr1* locus, where blocking *Commd1* transcription through the *Zrsr1* DMR abolished methylation (27), and at murine KvDMR1, where truncation of *Kcnq1* transcription prevented methylation (28). In humans, evidence remains indirect: mutations at the *KCNQ1* locus in Beckwith-Wiedemann syndrome suggest that loss of transcription across the KCNQ1OT1:TSS-DMR prevents establishment of maternal DNA methylation in the germline (29,30).

Examples in human somatic cells show that aberrant transcriptional read-through can drive DNA methylation changes that cause disease. Deletion of the *EPCAM* polyadenylation signal accounts for ∼10% of cases of hereditary non-polyposis colorectal cancer (HNPCC, Lynch syndrome) by inducing read-through into and methylation-mediated silencing of the downstream DNA repair gene *MSH2*, restricted to tissues with active *EPCAM* expression (31,32). A similar mechanism was observed in α-thalassemia, where an 18 kb deletion encompassing the 3′ end of *LUC7L* led to transcription across the adjacent *HBA2* gene and its silencing by DNA methylation; this effect was reproduced in differentiated but not undifferentiated mouse embryonic stem cells, suggesting additional signals are required for promoter methylation in somatic cells (33). In patients with cblC disorder of vitamin B12 metabolism, a splice acceptor site mutation in *PRDX1* caused exon skipping and loss of transcriptional termination, resulting in read-through across the *MMACHC* promoter and DNA methylation (34).

However, a direct mechanistic link between transcriptional activity and the induction of DNA methylation at imprinting control regions as well as the molecular mechanisms behind are missing. The central hypothesis of this study is that transcription initiated at the AS-SRO and traversing the PWS-SRO induces DNA methylation. Because human oocytes are ethically inaccessible, a human iPSC-based in vitro model to recapitulate this process was established. iPSCs were selected not only for their human origin and genetic tractability but also for their capacity to differentiate, a feature of relevance since DNA methylation establishment may require additional stimuli beyond transcriptional activity alone, reminiscent of the dynamic epigenetic remodelling during oocyte maturation. To test the hypothesis, the AS-SRO was replaced with a doxycycline-inducible promoter (pTRE), enabling controlled transcription through the PWS-SRO upon doxycycline treatment. This inducible system required two components: a transactivator protein responsive to doxycycline and a regulatable promoter bound by the transactivator. In the first step, the transactivator was integrated into the AAVS1 safe-harbor locus on chromosome 19 using CRISPR/Cas9 (35). In a second step, the 880 bp AS-SRO on chromosome 15 was replaced with an inducible promoter. Upon induction, DNA methylation at the PWS-SRO was analyzed by targeted deep amplicon bisulfite sequencing, and transcriptional changes were assessed by whole-transcriptome RNA sequencing.

## MATERIAL AND METHODS

### Cell lines and cultivation of human induced pluripotent stem cells

Parental cell line DSMZi017-A was modified to express rtTA transactivator protein as described in Haake et al. (35), resulting in cell line DSMZi017-A-1. Cell lines are registered at hPSCreg (https://hpscreg.eu). Human iPSCs were maintained in StemMACS™ iPS-Brew XF medium (Miltenyi Biotec) on Vitronectin XF–coated plates (10 μg/mL, STEMCELL Technologies) at 37 °C, 5% CO₂, and 21% O₂. Cells were passaged every 5–6 days at ∼70–80% confluency using Gentle Cell Dissociation Reagent (STEMCELL Technologies) and replated as clumps onto freshly coated plates in pre-warmed medium.

### CRISPR/Cas9 genome editing

Genome editing was performed using a ribonucleoprotein (RNP) complex composed of single guide RNA (sgRNA) and Cas9 nuclease (Alt-R S.p. Cas9 V3, Integrated DNA Technologies). sgRNA (10 μM stock) was mixed with Cas9 (61 μM stock) to final concentrations of 0.9 μM and 0.3 μM, respectively, in Dulbecco’s phosphate-buffered saline (DPBS, Gibco™), and incubated for 15 min at room temperature. Human iPSCs (70–80% confluency) were dissociated with Accutase (Merck) and resuspended in P3 Nucleofector™ Solution (Lonza). For each reaction, 0.7 × 10⁶ cells were combined with the preformed RNP, 1–1.5 μg linearized donor plasmid (AS-SRO exchange), and 0.45 μL electroporation enhancer (IDT). Electroporation was performed using the 4D-Nucleofector™ System (Lonza) with program CB-150, according to manufacturer’s instructions. Transfected cells were plated in StemMACS™ iPS-Brew XF medium (Miltenyi Biotec) supplemented with 10 μM Y-27632 (STEMCELL Technologies) to enhance post-nucleofection survival. GuideRNAs are listed in Supplementary Table S2.

### Low density seeding and colony isolation

Following nucleofection, cells were cultured until approximately 70% confluency before being dissociated with Accutase (5 min, 37 °C) to generate a single-cell suspension. Cells were counted using a Neubauer chamber and seeded at low density (0.01 × 10⁶, 0.005 × 10⁶, or 0.001 × 10⁶ cells per 10 cm Vitronectin XF-coated dish) in culture medium supplemented with CloneR™ (STEMCELL Technologies) to enhance single-cell survival. After 24 h, the medium was replaced with standard culture medium. Cells were cultured for up to 7 days until individual colonies became visible. Colonies with typical hiPSC morphology were manually isolated following brief incubation with Gentle Cell Dissociation Reagent (3 min, 37 °C) and transferred into Vitronectin XF-coated 96-well plates containing culture medium supplemented with 10 μM Y-27632 to enhance cell survival. After 24 h, cells were maintained in standard culture medium with daily medium changes. Colonies were expanded for 4–6 days before passaging into duplicate 96-well plates. One plate was used for direct lysis and genotyping, while the corresponding duplicate plate was maintained for further expansion of correctly targeted clones. Direct lysis of clonal cell lines was performed at 70–90% confluency using lysis buffer (GeneAmp buffer II, 0.5 mM MgCl_2_, 0.045 % Tween 20, 0.045 % NP-40, 50 μg/ml Proteinase K, incubation at 56 °C for 2 h followed by enzyme inactivation at 95 °C for 10 min. 1 μl of lysate was subsequently used for PCR-based genotyping.

### Trilineage directed differentiation

The differentiation potential of iPSCs was assessed using the StemMACS™ Trilineage Differentiation Kit (Miltenyi Biotec) following the manufacturer’s protocol. Briefly, cells were seeded onto Matrigel-coated plates and directed towards ectoderm, mesoderm, and endoderm lineages over 7 days. Differentiation efficiency was evaluated by flow cytometry using lineage-specific markers: PAX6 and SOX2 (ectoderm), CD184 and SOX17 (endoderm), and CD140b and CD144 (mesoderm). For DNA methylation analysis, cells were cultured in the presence or absence of 2.5 μg/mL doxycycline. Antibodies are listed in Supplementary table 1.

### Differentiation into hepatic-like cells

iPSCs were differentiated into hepatic-like cells (HLCs) using the STEMdiff™ Hepatocyte Kit (STEMCELL Technologies) according to the manufacturer’s protocol. Cells were seeded onto Laminin-521–coated plates (10 μg/mL, STEMCELL Technologies) and differentiated through three stages: definitive endoderm (days 1–5), hepatic progenitors (days 5–10), and maturation into HLCs (days 10–21). Cells were harvested at days 0, 5, 10, and 21 for DNA methylation analysis and whole transcriptome sequencing. Differentiation was performed with or without doxycycline (2.5 μg/mL), which was added 48 h prior to induction and maintained with each medium change. Successful differentiation was confirmed by morphology and transcriptomic analysis.

### Flow cytometry analysis

Flow cytometry was used to analyse the expression of surface and intracellular markers to assess pluripotency and differentiation status of iPSCs. Single-cell suspensions were prepared using Accutase (Merck). For surface staining, cells were incubated with fluorophore-conjugated antibodies (1:50) in DPBS for 30 min at 4 °C in the dark. For intracellular staining, cells were fixed in 4% paraformaldehyde for 30 min at 4 °C, permeabilized with 0.1 % Triton X-100 (Sigma), and stained with fluorophore-conjugated antibodies (1:50) as described. Unstained and isotype controls were included for gating. Data were acquired on a CytoFLEX S flow cytometer (Beckman Coulter) with 10,000 events per sample and analyzed using CytExpert software. Antibodies are listed in Supplementary Table S1.

### STR typing, mycoplasma detection and virus testing

Short tandem repeat (STR) typing, mycoplasma detection, and virus diagnostics were performed as a service by Leibniz Institute DSMZ GmbH (Germany).

### DNA methylation analysis by deep bisulfite amplicon sequencing

Genomic DNA was extracted using the QIAamp DNA Blood Mini Kit (Qiagen) and 500 ng per sample was bisulfite converted with the EZ DNA Methylation-Gold™ Kit (Zymo Research) following the manufacturer’s instructions. Target regions were amplified as described in (37), using primers from (38).

### Polymerase Chain Reaction – PCR

PCR amplification of plasmid DNA, genomic DNA, and bisulfite-converted DNA was performed using the HotStarTaq® Master Mix Kit (Qiagen) or the GoTaq® G2 Hot Start Green Master Mix Kit (Promega) according to the manufacturers’ instructions. Primers are listed in Supplementary Table S2.

### RNA isolation and quantitative RT-PCR

Total RNA was extracted from cell pellets using the RNeasy Plus Mini Kit with gDNA Eliminator columns (Qiagen), following the manufacturer’s instructions. For hepatic-like cells, RNA was isolated using the Direct-zol RNA Miniprep Plus Kit (Zymo Research) with TRI Reagent®–based lysis. RNA was eluted in RNase-free water. Total RNA was treated with RQ1 RNase-Free DNase (Promega) and reverse transcribed into cDNA using the LunaScript RT SuperMix Kit (New England Biolabs). Quantitative PCR was performed with SsoAdvanced™ Universal SYBR® Green Supermix (Bio-Rad) on an Applied Biosystems 7500 Fast Real-Time PCR System. Each 20 μL reaction contained 12.5 ng cDNA and 10 pmol of each primer. Cycling conditions were 95 °C for 2 min, followed by 40 cycles of 95 °C for 3 s and 60°C for 25 s. Primers are listed in Supplementary Table S2.

### Whole transcriptome sequencing and analysis

Whole transcriptome sequencing was performed on a NovaSeq 6000 instrument using paired-end reads of 150 bp by the sequencing facility GMAK, Helmholtz Zentrum für Infektionsforschung, Inhoffenstraße 7 in Braunschweig. Adapter and quality trimming of paired-end RNA-seq reads was performed using fastp v0.23.2 (39). Transcript abundance was quantified with Salmon v1.10.2 using sequence- and GC-bias correction and a custom GENCODE release 45 reference containing the inserted pTRE sequence (40). Transcript-level estimates were imported into R using tximport v1.28.0 (41), and differential gene expression was analyzed with DESeq2 v1.38.0 using internal size-factor normalization and Benjamini–Hochberg correction (42). Genes with |log2FC| ≥ 1 and adjusted p ≤ 0.05 were considered differentially expressed. Heatmaps and expression plots were generated using pheatmap v1.0.12 and gplots v3.2.0, respectively. For visualization of transcription across the modified PWS/AS region, reads were aligned to the modified GRCh38 reference using STAR v2.7.11b (43), processed with SAMtools v1.12, and visualized using Gviz in R (44). Coverage plots were used for qualitative assessment of transcript structure and were not normalized between samples. Gene Ontology enrichment analysis of differentially expressed genes was performed using clusterProfiler v4.14.0 (45).

## RESULTS

### Inducible iPSC in vitro model to drive transcription across the PWS-SRO

To engineer a human iPSC-based model, the AS-SRO element (880 bp) on chromosome 15 was replaced by a doxycycline-inducible promoter (pTRE, 376 bp), in iPSC line DSMZi017-A-1 (35), generating cell lines DSMZi017-A-2, DSMZi017-A-3, DSMZi07-A-4 and DSMZi017-A-5. The parental line DSMZi017-A was derived from a patient with Angelman syndrome carrying a large maternal deletion in 15q11–q13 (46). This deletion results in loss of the maternally methylated PWS/AS locus, thereby allowing precise targeting of the remaining, unmethylated paternal allele for investigation of DNA methylation establishment. The AS-SRO was replaced with a tetracycline-responsive promoter, allowing controlled transcription initiation by a transactivator protein (rtTA). Replacing only the AS-SRO maintained the natural orientation and spacing between the upstream regulatory element (AS-SRO) and the downstream element (PWS-SRO), as well as the surrounding genomic context. PCR screening was used to verify correct integration and the absence of the AS-SRO, using primer pairs outside the homology arms and inside the construct from both sides (Supplementary Fig. S2A). Additionally, a primer pair spanning the entire AS-SRO region was used to confirm pure cell populations with exchanged AS-SRO. After each round of engineering, a quality control panel was applied to confirm genomic integrity (not shown) and to ensure that the iPSCs maintained their pluripotency and differentiation potential (Supplementary Fig. S2B, S2C).

To test activation of the inducible transcript, RT-PCR was performed using multiple primer pairs covering upstream regions of the 35 kb interval between the AS-SRO and PWS-SRO (Fig. 1A). Amplified products were subjected to Sanger sequencing to deduce the splicing pattern (Fig. 1B). In clonal line DSMZi017-A-4, doxycycline treatment for 48 h induced robust expression of transcripts containing upstream exons, whereas control cells showed only low-level signals from the most upstream amplicons (primer pairs 1/2 and 3/4) (Fig. 1C). As a positive control, endogenous *SNRPN* expression was detected with primer pair 7/8 under both conditions, confirming RNA integrity and assay reliability. Amplicons generated with primer pairs 1/2, 3/4, and 5/6 corresponded to unspliced transcripts, co-linear with genomic DNA. By contrast, products amplified with primer pairs 1/8 and 2a/8 revealed splicing events, with transcripts initiating either at the inducible promoter or at the upstream exon U6.5 and splicing onto exon 2 of *SNURF-SNRPN* (Fig. 1B). These splicing patterns are consistent with those previously described for AS-SRO–initiated transcripts in human oocytes (13).

**Figure 1:**
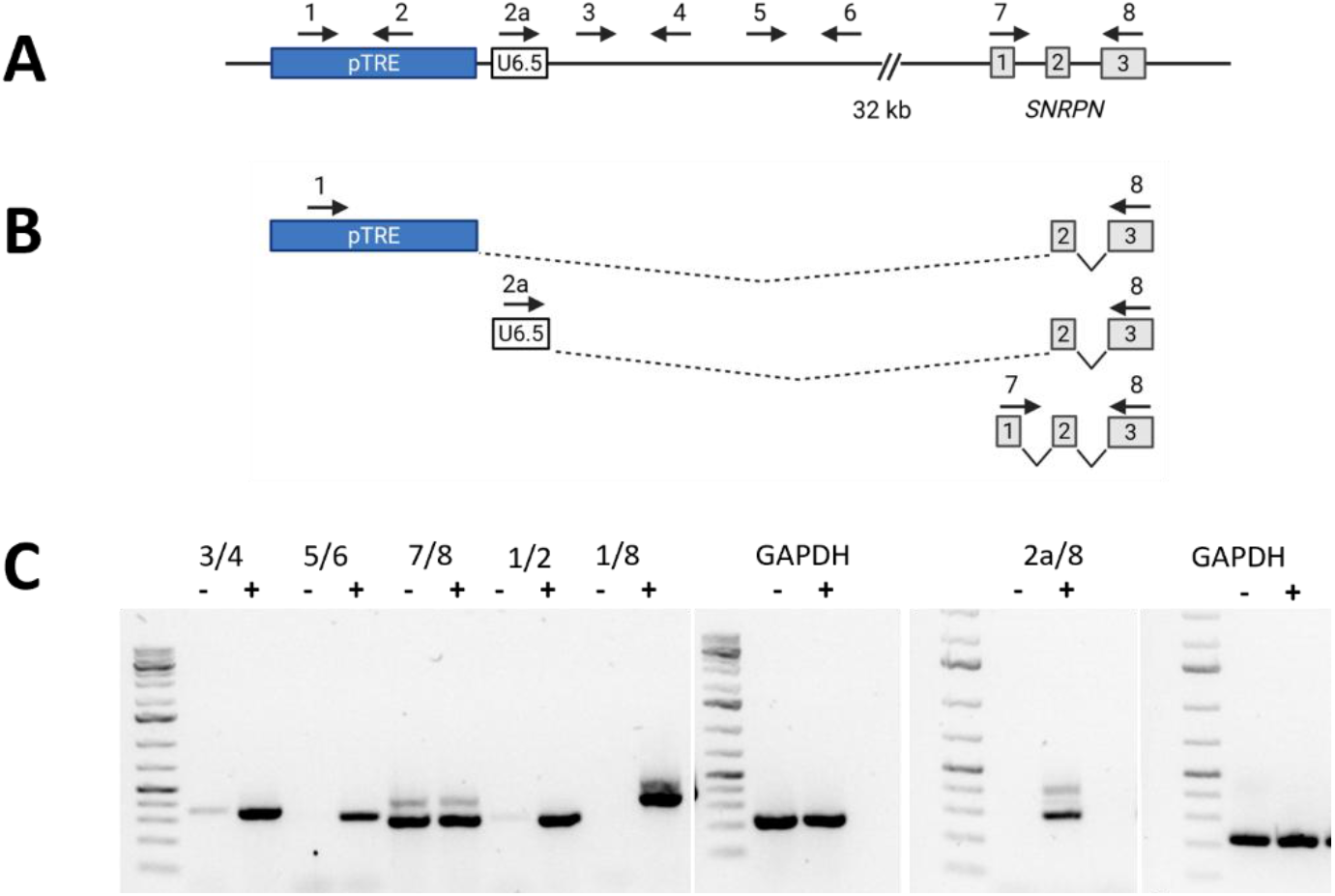
Inducible expression of the upstream transcript. (A) Schematic overview of the primer positions used for RT-PCR across the region surrounding the inducible promoter. The schematic is not drawn to scale. Blue: pTRE inducible promoter, inserted at the position of the original AS-SRO. (B) Splice variants reconstructed after Sanger sequencing of RT-PCR products. (C) Gel electrophoresis of RT-PCR products obtained from untreated (–) and doxycycline-induced (+) DSMZi017-A-4 cells after 48 hours. Only *SNRPN* (primers 7/8) and *GAPDH* were expressed in both conditions at similar intensities, serving as positive controls for RNA quality and cDNA synthesis. –RT controls (without reverse transcriptase) were negative (data not shown). Illustrations were made by BioRender.

### Acquisition of DNA methylation after induced transcription

After stable induction of transcription across the PWS-SRO, DNA methylation establishment was analyzed by deep bisulfite amplicon sequencing (37) in three clonal cell lines (DSMZi017-A-2/-4/-5). The amplified region (spanning 240 bp) is located approximately 33.5 kb downstream of the inducible promoter within the PWS-SRO region, including *SNRPN* exon 1 and at the beginning of the CpG island 77 (Supplementary Fig. S1B). Acquisition of DNA methylation was determined in undifferentiated iPSCs (Supplementary Fig. S3A, B), undirected differentiation to embryoid bodies (EB) (Supplementary Fig. S3C, D) and short-term directed differentiation (trilineage differentiation) into derivatives of the three germ layers (Fig. 2). Although expression of the induced transcript could be observed in all experiments by RT-qPCR, a gain of DNA methylation at the PWS-SRO was only achieved in the endoderm lineage upon directed trilineage differentiation, showing up to 10% DNA methylation (Fig. 2, Supplementary Fig. S4A). Induction of the transcript 48 hours before differentiation did not yield a higher level of DNA methylation (Supplementary Fig. S4B). Because cell morphology started to deteriorate from day 5 on, doxycycline was withdrawn at day 4. But no gain of DNA methylation was observed, indicating that presence of the induced transcript between days 5 to 7 is essential (Supplementary Fig. S4C).

**Figure 2:**
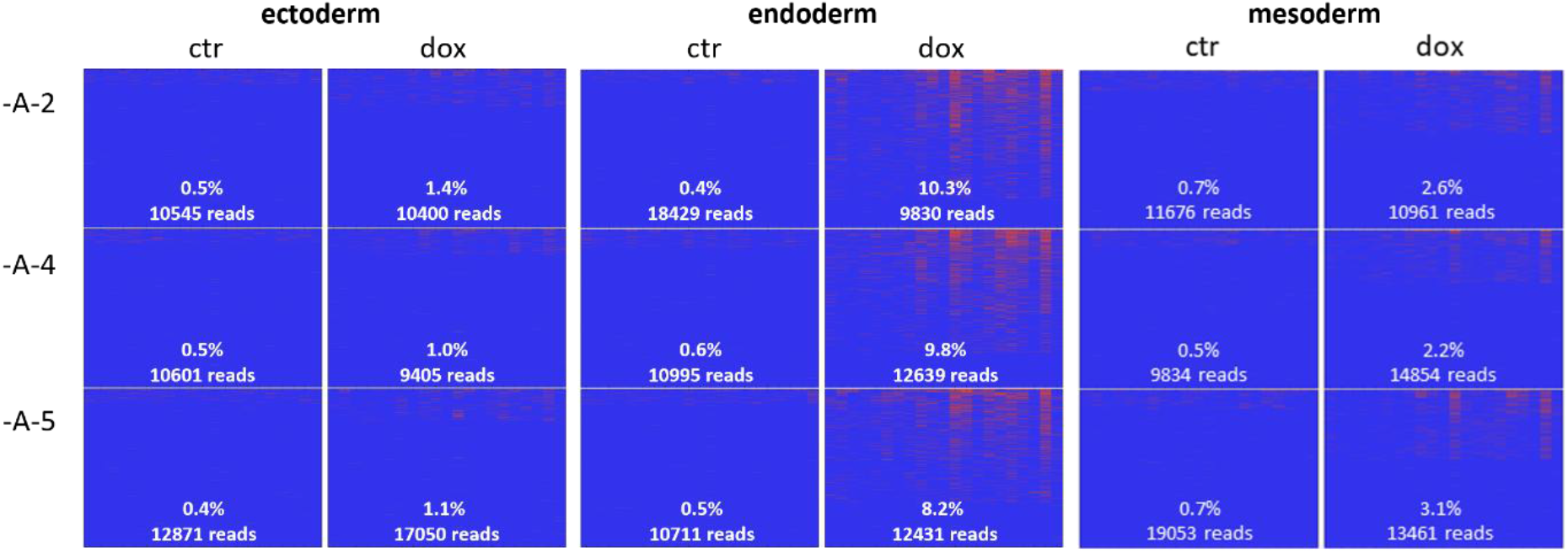
DNA methylation analysis after differentiation into ecto-, endo- and mesoderm. Heatmaps showing results of bisulfite sequencing of the *SNRPN* amplicon. CpGs are in columns, reads are in rows, blue indicates non-methylated CpGs, red indicates methylated CpGs. Percentage of overall DNA methylation and number of reads are given in the plots.

Since DNA methylation establishment at the PWS-SRO was only observed in endodermal cells derived from trilineage differentiation but reached relatively low levels (up to 10%), directed differentiation into definitive endoderm in 5 days was performed. This approach aimed to enrich a more homogeneous and transcriptionally committed endodermal population, which could potentially support higher levels of DNA methylation upon induction of the inducible transcript. Successful differentiation and presence of the induced transcript was confirmed (Supplementary Fig. S5A, B). Acquisition of DNA methylation was reliably observed up to a level of about 5% (Supplementary Fig. S5C) confirming the permissive state of endodermal cells but indicating that a period of five days of differentiation might be too short for effective DNA methylation.

After observing a gain of DNA methylation between 5% and 10% in the endodermal lineage, hepatic differentiation was applied to extend the differentiation period to 21 days and promote the development of more mature endodermal derivatives. Since hepatic-like cells originate from the endoderm, iPSCs were differentiated using a three-stage hepatic differentiation protocol including the generation of definitive endoderm over a period of five days, further maturing into hepatic progenitor cells (day 10) and differentiation into hepatic-like cells, continuing until day 21. Cells were harvested at days 5, 10 and 21 for analysis by RNA-seq and amplicon bisulfite sequencing. Whole transcriptome RNA-seq confirmed successful differentiation into the endodermal lineage and hepatic specification and induction of transcription initiating at the inducible promoter at the AS-SRO position (Supplementary Fig. S6A). However, transcription at the inducible promoter ceased from after day 5 and was barely observed at days 10 and 21 (Supplementary Fig. S6B). Since GFP expression also declined at days 10 and 21 (Supplementary Fig. S6C) and both transcripts were under the control of an rtTA-inducible promoter, silencing of rtTA transactivator protein expression during ongoing differentiation towards hepatic-like cells was assumed. DNA methylation changes were analysed during hepatic differentiation in undifferentiated iPSCs (day 0), definitive endoderm (day 5), hepatic progenitor cells (day 10) and hepatic-like cells (day 21) (Fig. 3). Cells treated with doxycycline showed a notable increase in DNA methylation of up to 10% on day 5 of differentiation. DNA methylation levels stabilized or slightly increased by day 10 and remained relatively stable on day 21 in clonal line DSMZi017-A-2 (10.6%), while a decrease was observed in DSMZi017-A-4 (6.4%) and DSMZi017-A-5 (3.3%). In addition to changes in overall DNA methylation levels, the pattern of methylated CpGs within the amplicon changed from single, rather highly methylated CpGs to fully methylated reads. On day 5 of differentiation, individual sequencing reads displayed partial methylation, affecting only a few specific CpG sites, rather than the entire read. By day 10 and continuing through day 21, individual reads more frequently exhibited methylation across nearly all 21 CpG sites, indicating a shift from partial to global methylation within single molecules. Although the number of methylated reads slightly decreased at later time points, the reads that were methylated tended to show dense, continuous methylation, suggesting a stabilization and expansion of the methylation pattern over time. In summary, acquisition of DNA methylation at the PWS-SRO upon induction of transcription is robust and reproducible in the developed system, demonstrating for the first time a direct connection between transcription and DNA methylation establishment in a human system.

**Figure 3:**
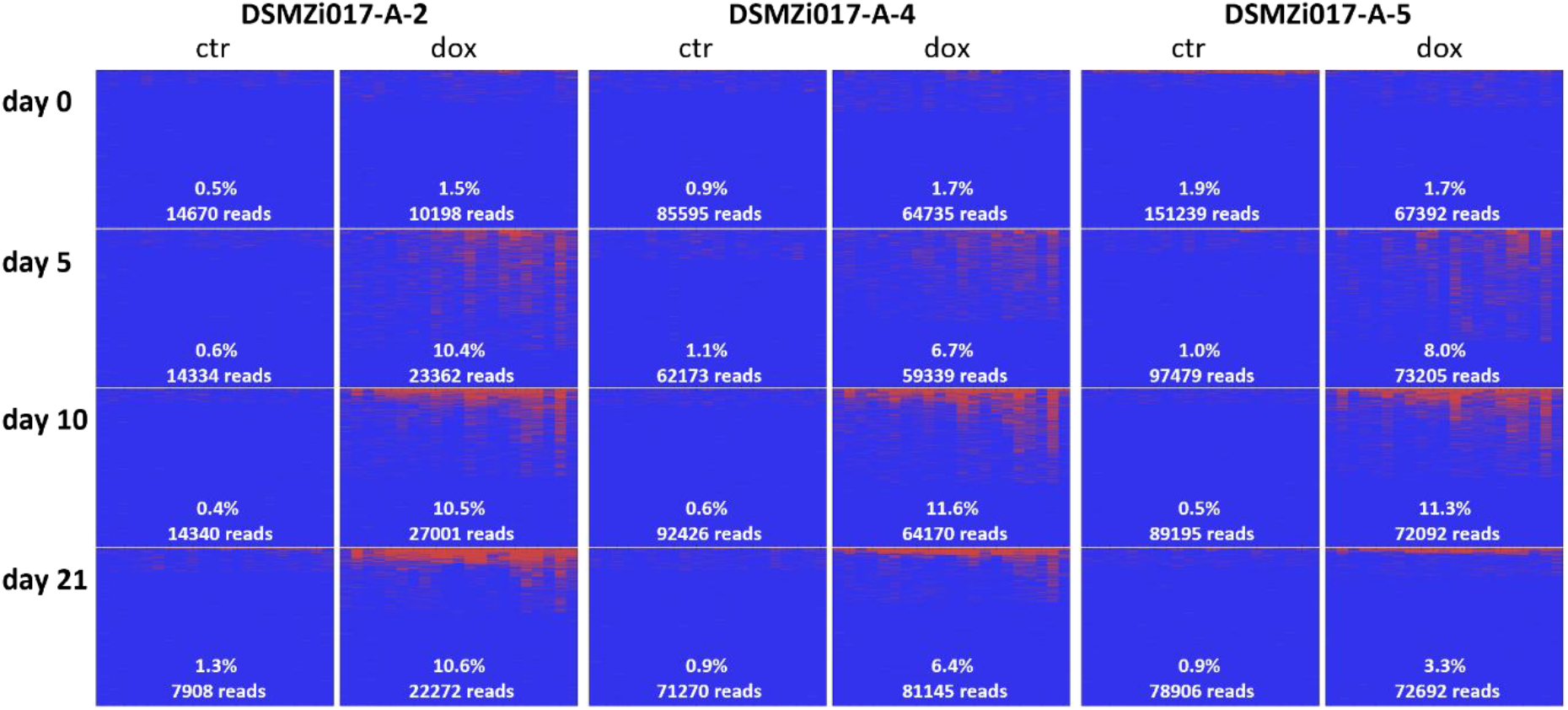
DNA methylation analysis in hepatic-like cells. Heatmaps showing results of bisulfite sequencing of the *SNRPN* amplicon at day 0 (iPSCs), day 5 (definitive endoderm), day 10 (hepatic progenitor cells), and day 21 (hepatic-like cells), from top to bottom. CpG sites are shown in columns; sequencing reads are in rows. Blue indicates unmethylated CpGs; red indicates methylated CpGs. The overall percentage of DNA methylation and the number of reads are indicated in each plot.

### Impact of the induced transcript on gene expression at the PWS/AS locus

To determine the overall influence of doxycycline treatment on gene expression in iPSCs, differential gene expression analysis was performed comparing undifferentiated iPSCs not induced or induced by doxycycline for 10 days. Gene expression was highly similar between the two groups, only two genes were found to be downregulated, *SNRPN* and *BIVM-ERCC5*, whereas several genes were upregulated in doxycycline-treated samples (Fig. 4A). Downregulation of *SNRPN* indicates that its expression indeed is influenced by the induced transcript, whereas the mechanism of regulation of *BIVM-ERCC5* (located on chromosome 13) by doxycycline or the induced transcript remains elusive. Gene ontology analysis of DEGs showed enrichment of processes involved in mesodermal tissue development, indicating that doxycycline treatment of iPSCs per se would favour differentiation into that direction (Supplementary Fig. S7A). Downregulation of *SNRPN* as a consequence of the induced transcript extending across the PWS-SRO, which encompasses the promoter of *SNRPN*, was further assessed by RT-qPCR in undifferentiated iPSCs, embryoid bodies at d21 and in definitive endoderm at d5 (Fig. 4B). In undifferentiated iPSCs, doxycycline treatment led to a significant downregulation of *SNRPN* compared to untreated controls. In contrast, embryoid body (EB) differentiation for 21 days did not result in significant changes in *SNRPN* levels upon induction of the transcript. During differentiation to definitive endoderm, *SNRPN* expression was significantly reduced in all three clonal lines on day 5 following doxycycline treatment. Following trilineage differentiation, *SNRPN* expression was assessed on day 7 (Fig. 4C). In the ectodermal lineage, *SNRPN* was moderately increased. The observed upregulation of *SNRPN* during ectodermal differentiation may be attributed to the activation of the long non-coding RNA *SNHG14*, which includes *SNRPN* exons and is specifically expressed in neuronal cells, a lineage derived from the ectoderm (47). In the endodermal lineage, *SNRPN* expression was significantly reduced in doxycycline-treated samples compared to controls. Although only one clone was available for the trilineage endoderm analysis, the observed reduction was consistent with the independent day 5 endoderm differentiation. In contrast, mesodermal differentiation did not result in significant differences in *SNRPN* expression between treated and untreated cells. When doxycycline was withdrawn at day 4, resulting in markedly reduced transcription across the PWS-SRO, no significant differences in *SNRPN* expression were detected between induced and control conditions in any of the three germ layers on day 7 (Supplementary Fig. 7B). Together, these results demonstrate that expression of the induced transcript is associated with reduced *SNRPN* expression in undifferentiated iPSCs and during endodermal differentiation, while no consistent effect was observed in mesodermal or ectodermal conditions. *SNRPN* expression during hepatic differentiation was assessed in the RNA-seq dataset (Fig. 4D). Consistent with the RT-qPCR results, *SNRPN* expression was reduced upon induction in undifferentiated iPSCs (day 0), but no difference was detected at the later time points, coinciding with the loss of detectable inducible transcript expression. Since DNA methylation at the PWS-SRO regulates expression of several genes (*MKRN3*, *MAGEL2*, *NDN*, *NPAP1*, *SNURF/SNRPN*, snoRD RNA clusters, *SNHG14*) at the PWS/AS locus, normalized counts of protein-coding genes were analyzed (Fig. 4E). The downstream gene *UBE3A*, as well as the upstream genes *MAGEL2*, *NDN*, and *NPAP1*, showed no significant changes in expression, when comparing control and doxycycline treated cells. *MKRN3* was not expressed in iPSCs.

**Figure 4:**
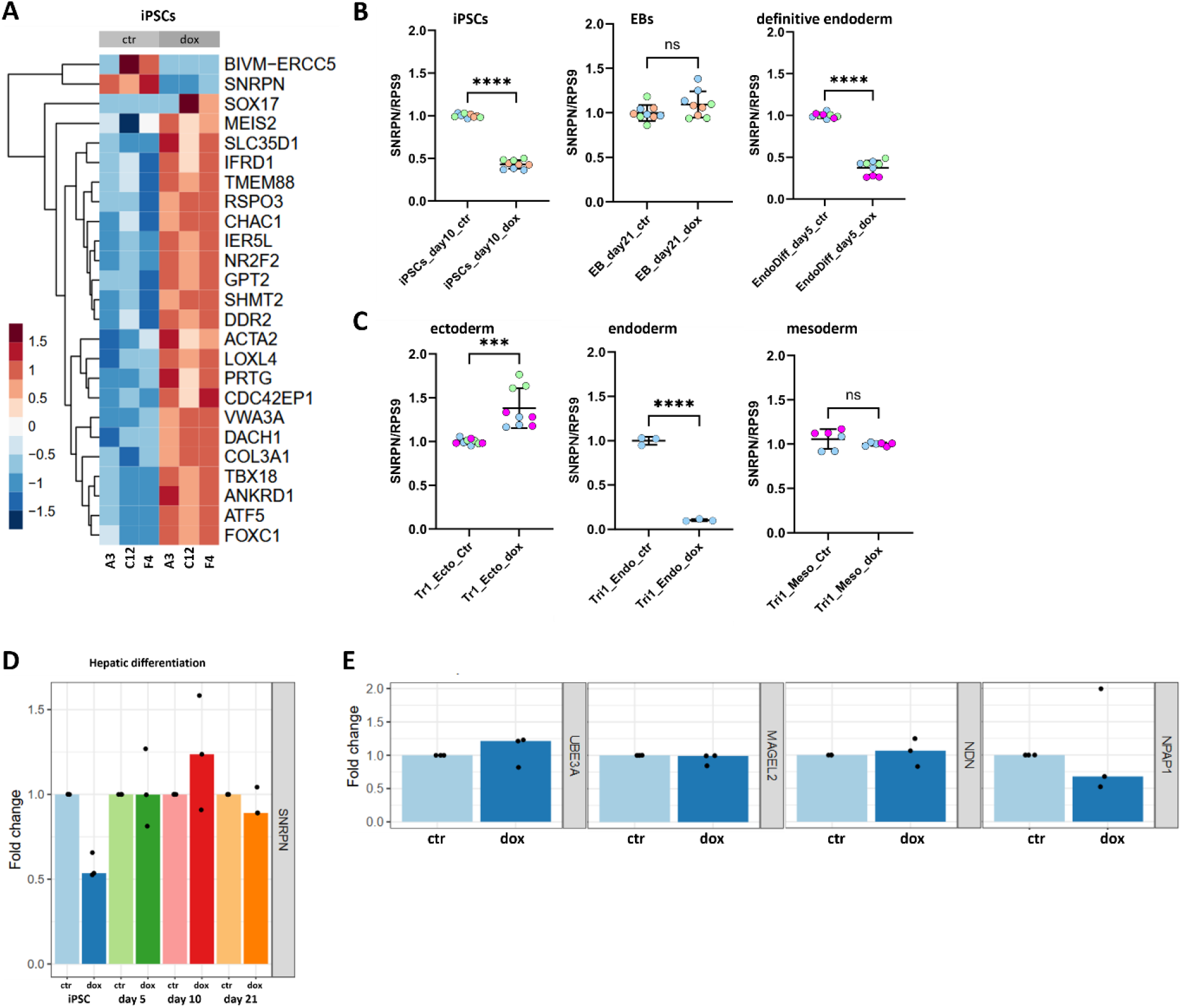
Gene expression analysis in doxycycline treated cells. (A) Differential gene expression analysis in undifferentiated iPSCs (DESeq2, clonal lines DSMZi017-A-2/-4/-5, -/+ doxycycline). Shown are top 25 genes (adjusted p-value ≤ 0.05, absolute log2(FC) ≥ 1, red indicating upregulation and blue indicating downregulation relative to the overall mean. (B) *SNRPN* expression in undifferentiated iPSCs, EB differentiation (d21), and differentiation to definitive endoderm (d5). Clonal lines DSMZi017-A-2: blue, -A-3: orange, -A-4: green. Gene expression was normalized to *RPS9* and calibrated to non-induced iPSCs. Error bars represent standard deviation. Statistical significance was determined using Student’s t-test; **** p ≤ 0.0001, n.s. = not significant. (C) *SNRPN* expression in derivatives of the three germ layers (trilineage differentiation) on day 7 with induction start on day 0 of differentiation. Clonal lines DSMZi017-A-2: blue, -A-4: green, -A-5: pink. Gene expression was normalized to *RPS9* and calibrated to non-induced iPSCs. Error bars represent standard deviation. Statistical significance was determined using Student’s t-test; **** p ≤ 0.0001, *** p ≤ 0.001, n.s. = not significant. (D) Fold change in *SNRPN* expression during hepatic differentiation, based on RNA-seq data. *SNRPN* expression in doxycycline-treated cells (dark bars) was compared to untreated control cells (light bars). Blue: iPSCs (day 0), green: definitive endoderm (day 5), red: hepatic progenitors (day 10), orange: hepatic-like cells (day 21). Shown are triplicate values and the mean. (E) Expression fold change of *UBE3A*, *MAGEL2*, *NDN*, and *NPAP1* showed no differences in expression between control and doxycycline-treated iPSCs.

## DISCUSSION

Despite growing evidence that transcription directs DNA methylation establishment at imprinted loci, the molecular mechanisms governing transcription-dependent de novo methylation remain poorly understood. This study provides the first direct evidence in a human system that transcriptional read-through initiates DNA methylation at an imprinting control region. The engineered hiPSC model enabled inducible transcription across the PWS-SRO, providing a tractable system to dissect the mechanistic link between transcription and de novo DNA methylation. The induced transcripts were spliced to *SNRPN* exon 2, thereby recapitulating a key structural feature of the endogenous oocyte-derived transcripts originating from the AS-SRO (13).

Gain of DNA methylation at the PWS-SRO was observed exclusively during directed differentiation into the endodermal lineage, emphasizing the importance of cellular remodelling processes for imprint establishment by DNA methylation. Previous models in human somatic cells or mouse undifferentiated ES cells failed to model the establishment of DNA methylation by transcription although transcriptional read-through across a CpG island was successfully induced (33,48,49). The absence of methylation in undifferentiated pluripotent cells and differentiated somatic cells, despite transcriptional activation, suggests that transcription is not sufficient by itself. Differentiation of PSCs is accompanied by extensive epigenetic remodelling, including dynamic changes in DNA methylation. Genome-wide studies have shown that CpG methylation patterns are significantly restructured during the transition from pluripotency to differentiated states (50,51). These changes may establish a chromatin context or provide cofactors necessary for the recruitment and function of de novo DNA methyltransferases. Transcription-dependent de novo methylation has also been described in differentiated human tissues, including α-thalassemia, Lynch syndrome and cblC deficiency (31,33,34). However, in these disorders it remains unclear whether methylation is established within mature somatic cells or during earlier developmental stages characterized by greater epigenetic plasticity. However, transcription and differentiation in general are not sufficient to induce DNA methylation. Differentiation needs to result in setting up a lineage-specific environment providing necessary cellular factors, as shown by establishment of DNA methylation only in the endodermal lineage. In contrast, mesodermal and ectodermal differentiation led to only low gain in methylation. During oogenesis, imprint establishment depends on a coordinated network of factors including DNMT3A, DNMT3L, SETD2, KDM1B and ZFP57 (26,52–56). Interestingly, several of these factors were also expressed during endodermal differentiation but were largely absent from mesodermal and ectodermal derivatives (Supplementary Fig. S8). Although expression alone does not demonstrate functional involvement, these observations suggest that endoderm transiently provides an epigenetic environment permissive for transcription-dependent de novo methylation. Interestingly, RNA sequencing also revealed elevated expression of NLRP7 in endoderm, a maternal-effect gene required for the establishment of maternal imprints (57–59). Whether its expression contributes to the permissive environment observed during endodermal differentiation remains an intriguing question for future studies.

The main limitation of the study are the consistently low levels (∼10 %) of DNA methlyation achieved by induction of transcription across the PWS-SRO throughout endodermal differentiation. Because of this low levels of DNA methylation, analysis of downstream processes like histone modifications or recruitment of DNA methyltransferases was not possible. Reduced *SNRPN* expression following induction of the transgene was consistently observed in undifferentiated iPSCs and endodermal cells, suggesting that activation of the inducible transcript negatively affects endogenous *SNRPN* transcription. This observation is consistent with transcriptional interference, whereby transcriptional activity of one locus negatively influences a neighbouring transcriptional unit in cis (60). In the present model, the inducible promoter is positioned upstream of the endogenous *SNRPN* promoter in tandem orientation. Tandem transcriptional interference has been shown to repress downstream promoters through mechanisms like promoter occlusion, premature termination, and transcription-associated chromatin remodelling (61,62). Transcriptional interference between tandem promoters depends on strength of promotors and is especially effective if the upstream promoter is stronger than the downstream promoter, which is read through (49). In the setting of this study, transcription initiation and elongation observed from the inducible promoter is substantially weaker than transcription from the *SNRPN* promoter. Active transcription from the *SNRPN* promoter may impede efficient read-through of the inducible transcript, thereby restricting the transcription-dependent establishment of DNA methylation. Importantly, *SNRPN* is not expressed during oogenesis, where maternal methylation at the PWS-SRO is established through transcription initiated from the upstream AS-SRO promoter (13). In contrast, *SNRPN* remained actively expressed in iPSCs and all germ layer derivatives examined in this study. Continuous loading of RNA polymerase II at the *SNRPN* promoter may therefore restrict efficient upstream read-through, limiting DNA methylation to a subset of approximately 10% of cells in which cellular conditions happen to be favorable by chance. To test this hypothesis, iPSCs were differentiated into hepatic progenitors and hepatic-like cells, as *SNRPN* is expressed at relatively low levels in liver tissue (Human Protein Atlas; https://www.proteinatlas.org/ENSG00000128739-SNRPN/tissue#rna_expression, accessed 06 September 2026). RNA sequencing confirmed reduced *SNRPN* expression during hepatic differentiation. However, the inducible transcript was no longer detectable presumably because of silencing of the rtTA transgene at the AAVS1 locus, precluding assessment of whether reduced *SNRPN* transcription could enhance DNA methylation establishment. Future studies will therefore require strategies that maintain stable transgene expression through hepatic differentiation to directly determine whether *SNRPN* transcription limits transcription-dependent methylation at the PWS-SRO.

An additional insight emerging from this study concerns the dynamics of de novo DNA methylation at the PWS-SRO during hepatic differentiation. Bisulfite sequencing revealed heterogeneous methylation patterns across individual DNA molecules, with some reads initially exhibiting methylation at only a few CpG sites before either progressing towards more extensive methylation or reverting to an unmethylated state. These observations suggest that DNA methylation at the PWS-SRO may occur as a progressive rather than an all-or-none process.

Despite the discussed limitations of consistent low level DNA methylation, probably related to *SNRPN* expression strength, the inducible human iPSC model provides a first experimental framework to investigate transcription-dependent DNA methylation and its temporal dynamics. Optimization of the model might allow dissection of molecular mechanisms underlying imprint establishment, questions that remain difficult to address in vivo.

## Supporting information

Haake_supplementary_material

## ACKNOWLEDGEMENTS

We thank Maren Kaufmann for excellent technical support and all members of the Human and Animal Cell Lines department for technical support and fruitful discussions.

## AUTHOR CONTRIBUTIONS

Josephine Haake: Conceptualization, Data curation, Formal analysis, Investigation, Validation, Visualization, Writing – original draft, Writing – review & editing. Alexander Lederhof: Methodology. Claudia Pommerenke: Data analysis. Laura Steenpass: Conceptualization, Funding acquisition, Project administration, Resources, Supervision, Writing – review & editing.

## SUPPLEMENTARY DATA

Supplementary Data are provided in file “Haake_supplementary_material”.

## DATA AVAILABILITY

Genetically modified cell lines generated and used in this study are registered in hPSCreg and available upon request. RNA sequencing data are available on request and will be deposited in ArrayExpress at ENA.

