## Supplementary material for "Transcription-driven establishment of DNA methylation at the PWS/AS imprinted region": Haake_supplementary_material

**Supplementary methods**

**Immunofluorescence staining**
Immunocytochemistry was performed to assess nuclear stem cell marker expression in iPSCs. Cells were fixed with 4% paraformaldehyde (Thermo Scientific) for 15 min at room temperature, permeabilized and blocked in 5% milk powder with 0.4% Triton X-100, and incubated with primary antibodies (1:200) for 1 h. After washing, cells were incubated with fluorophore-conjugated secondary antibodies (1:1,000) for 30 min in the dark, counterstained with DAPI (5 µg/mL, Sigma-Aldrich), and imaged using a Zeiss Axio Observer fluorescence microscope with ZEN 3.3 software. Representative images were acquired under identical exposure settings. Antibodies are listed in Supplementary Table S1.

**Embryoid body (EB) formation and undirected differentiation**
Embryoid bodies were generated by dissociating iPSCs into single cells using Accutase (Merck) and seeding 1.2 × $10^4$ cells per well in low-adhesion, round-bottom 96-well plates (Nunclon™ Sphera, Thermo Scientific) in AggreWell™ EB Formation Medium supplemented with Y-27632 (10 µM, STEMCELL Technologies). After 24 h, medium was replaced with EB medium, which was subsequently changed every other day. After 7 days, EBs were transferred to gelatin-coated 24-well plates (0.2% gelatin) for spontaneous differentiation. Four EBs were plated per well in EB medium, with half-medium changes performed daily. For DNA methylation analysis, cells were differentiated in the presence or absence of doxycycline (2.5 µg/mL), which was added from day 0 and refreshed with every medium change.

**Definitive endoderm differentiation**
Differentiation of iPSCs into definitive endoderm was performed using the STEMdiff™ Definitive Endoderm Kit (STEMCELL Technologies) according to the manufacturer's instructions. For DNA methylation analysis, cells were cultured in the presence or absence of 2.5 µg/mL doxycycline, which was added 48 h prior to differentiation and maintained with every medium change. On day 5, cells were harvested for DNA methylation and RNA expression analyses, and endoderm identity was confirmed by flow cytometry using CD184 and SOX17 as markers.

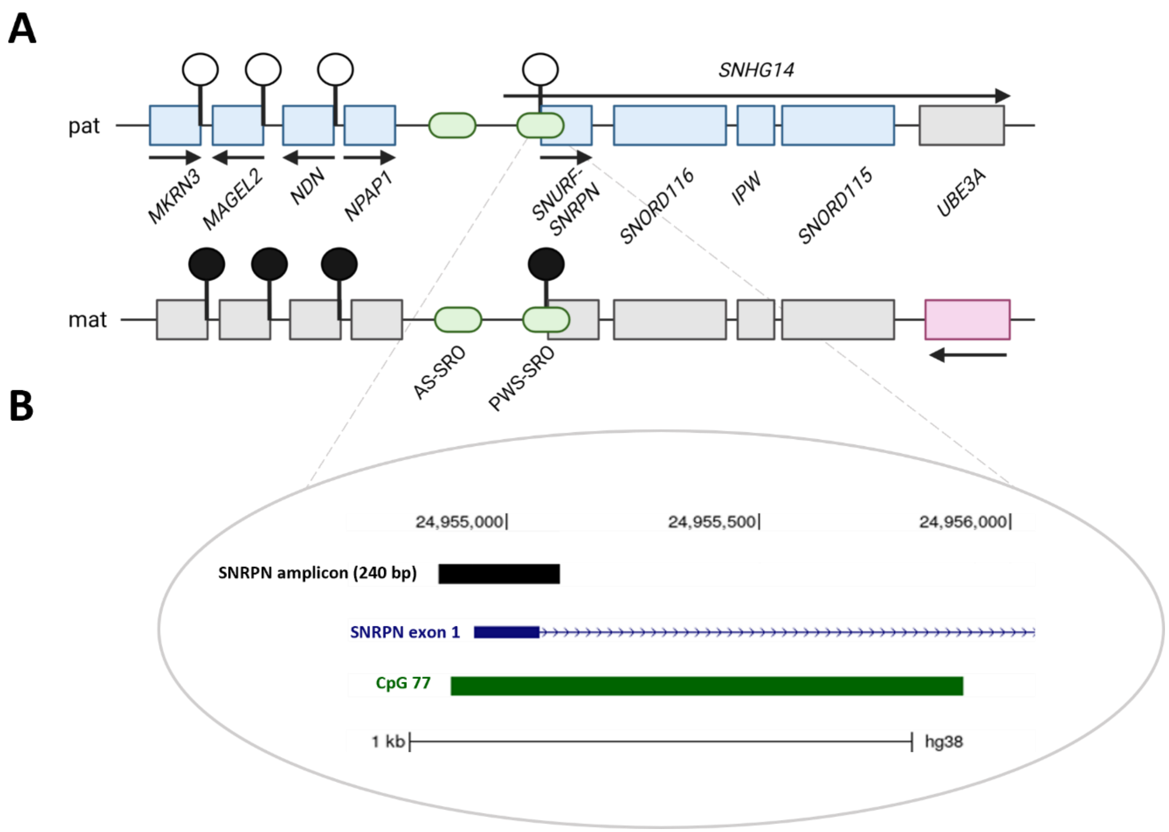

**Supplementary Figure S1. The Prader-Willi / Angelman syndrome locus on human chromosome 15.**
**(A)** Schematic view of the PWS/AS locus, not drawn to scale. The paternal allele is shown at the top, maternal allele shown below. Gene expression status as observed in the brain. Blue: Genes expressed from the paternal allele, red: genes expressed from the maternal allele, green: bipartite imprint control region consisting of PWS-SRO and AS-SRO, black lollipop: methylated region, white lollipop: not methylated region. Arrows indicate direction of transcription.
**(B)** Zoom-in into the 5'-end of the PWS-SRO. Green: CpG island CpG77, blue: *SNRPN* exon 1 with orientation of transcription, black: 240 basepair amplicon analyzed for DNA methylation by deep amplicon sequencing. All elements are drawn to scale based on the UCSC Genome Browser (GRCh38/hg38).

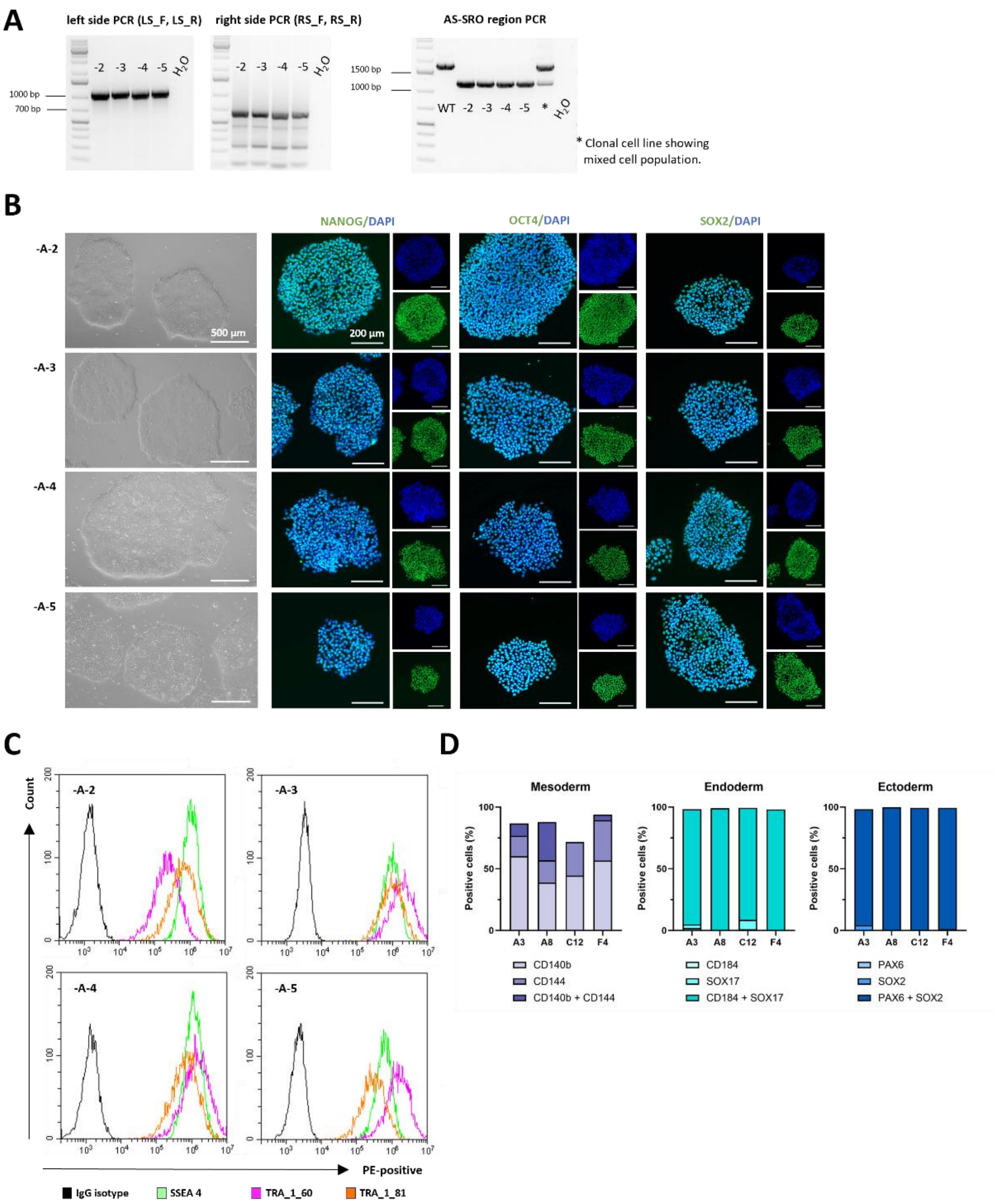

**Supplementary Figure S2. Characterization of clonal iPSC lines after CRISPR/Cas9-mediated AS-SRO exchange in clonal lines DSMZi017-A-2/-A-3/-A-4/-A-5.**

**(A)** To confirm correct exchange of the AS-SRO by the inducible promoter, PCR was performed using primers spanning from outside the homology arms into the inducible promoter, yielding a 1130 bp fragment (left side) and a 717 bp fragment (right side) in correctly targeted clones. In addition, the entire integration site (AS-SRO region) was amplified. A 1194 bp fragment was detected in clonal lines DSMZi017-A-2/3/4/5, confirming a clean cell population.

**(B)** Stem cell morphology and expression of nuclear pluripotency markers was confirmed by brightfield and fluorescent microscopy. Nuclear stem cell markers NANOG, OCT4 (POU5F1) and SOX2 in green, nuclei counterstained with DAPI (blue). Scale bars 500 µm and 200 µm.

**(C)** Flow cytometry of surface pluripotency markers confirmed expression of SSEA4 (green), TRA-1-60 (pink), and TRA-1-81 (orange).

**(D)** Engineered iPSC lines were differentiated towards ectoderm, endoderm, and mesoderm lineages, and analyzed by flow cytometry after 7 days using two lineage-specific markers per germ layer. The majority of cells were double-positive for ectoderm (blue) and endoderm (turquoise) markers, while mesoderm-differentiated cells (purple) were predominantly single-positive, reflecting successful but heterogeneous lineage commitment.

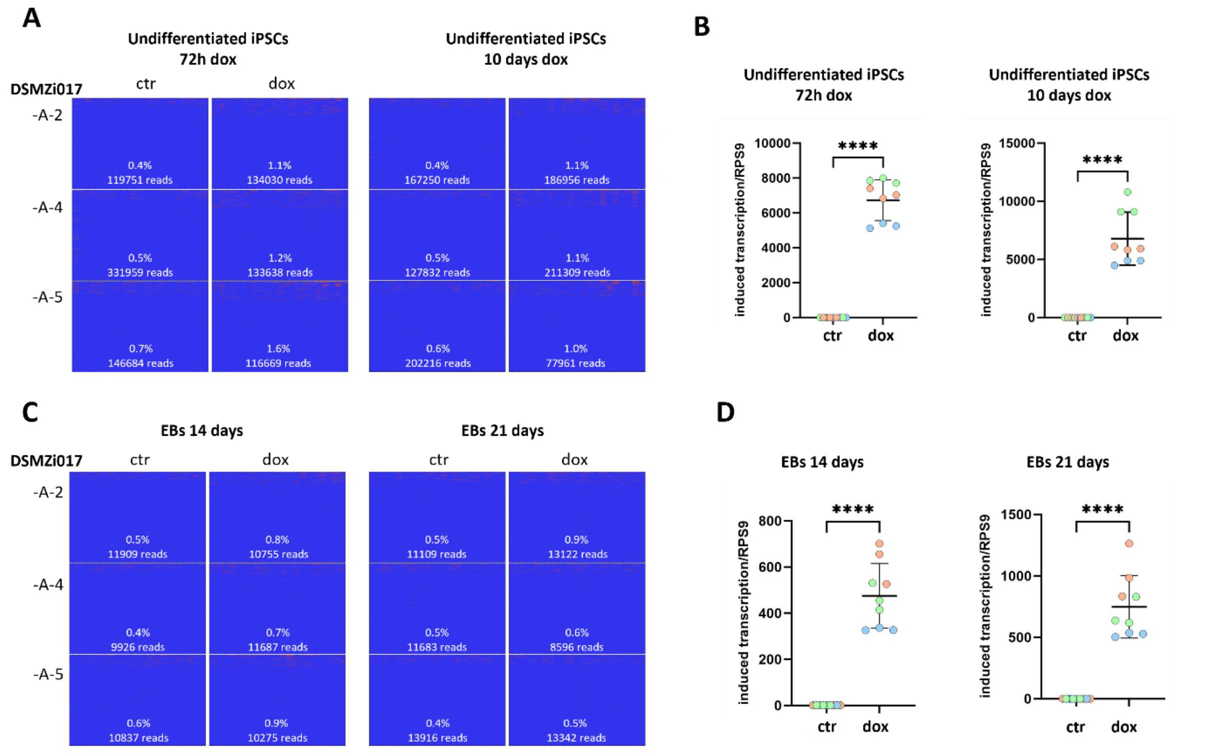

**Supplementary Figure S3. DNA methylation analysis and induction of transcription in undifferentiated iPSCs and embryoid bodies (EBs) in clonal lines DSMZi017-A-2/-A-4/-A-5.**

**(A)** Heatmaps of bisulfite sequencing results for the analyzed amplicon in undifferentiated iPSCs 72h and 10 days after induction of the induced transcript. CpGs are in columns, reads are in rows, blue indicates non-methylated CpGs, red indicates methylated CpGs. Percentage of overall DNA-methylation and number of reads are given in the plots.

**B)** RT-qPCR in undifferentiated iPSCs 72h and 10 days after induction confirming induction of the transcript across the PWS-SRO, with significantly higher expression in doxycycline-treated vs. untreated cells (Student's t-test; ****$p \leq 0.0001$). Expression was normalized to *RPS9* and calibrated to non-induced iPSCs. Values of technical triplicates are shown: blue: -A-2; orange: -A-3; green: -A-4.

**(C)** Heatmaps of bisulfite sequencing results for the analyzed amplicon in EBs on day 14 and day 21 of differentiation. CpGs are in columns, reads are in rows, blue indicates non-methylated CpGs, red indicates methylated CpGs. Percentage of overall DNA-methylation and number of reads are given in the plots.

**(D)** RT-qPCR confirming induction of the transcript across the PWS-SRO in EBs on day 14 and day 21 of differentiation, with significantly higher expression in doxycycline-treated vs. untreated cells (Student's t-test; ****$p \leq 0.0001$). Expression was normalized to *RPS9* and calibrated to non-induced iPSCs. Values of technical triplicates are shown: blue: -A-2; orange: -A-3; green: -A-4.

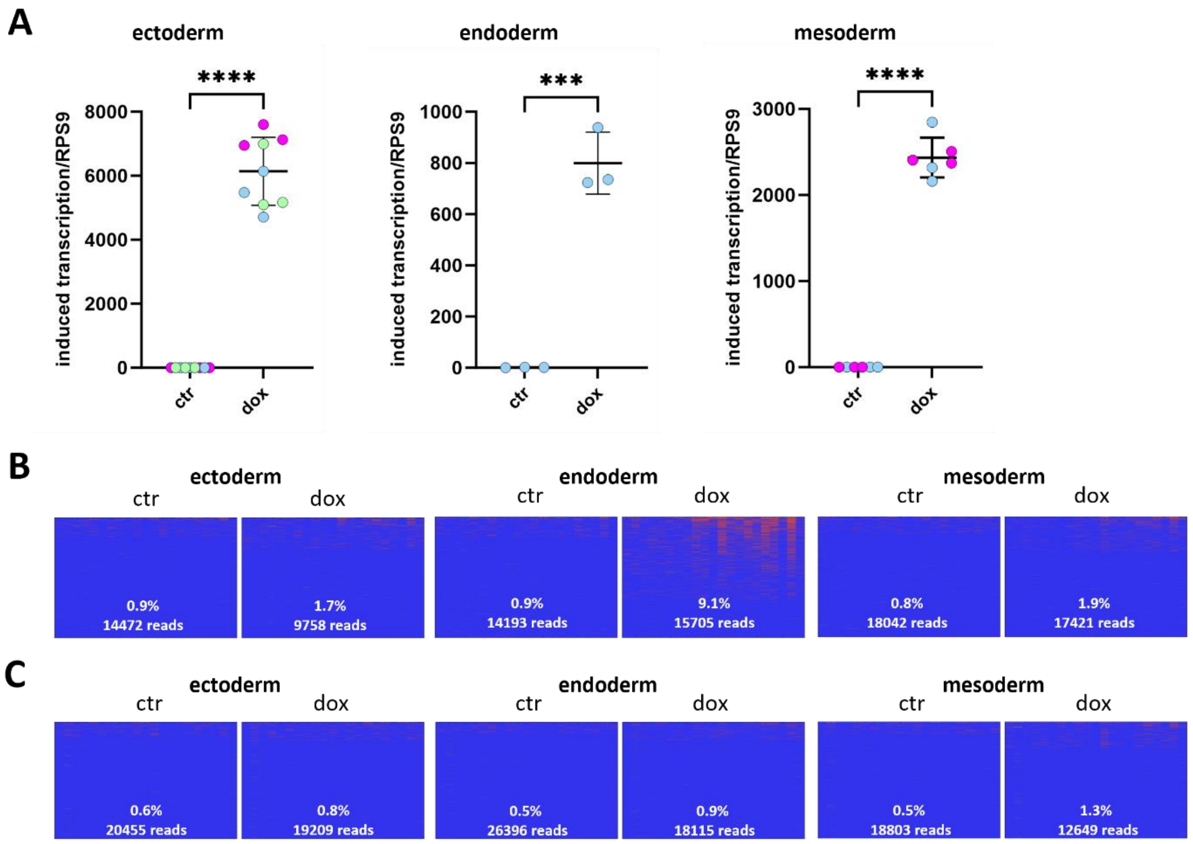

**Supplementary Figure S4. Expression of the induced transcript and DNA methylation at d7 of trilineage differentiation.**

**(A)** RT-qPCR confirming induction of the induced transcript (on day 0) measured on day 7 of trilineage differentiation. Results for ectoderm, endoderm, and mesoderm are normalized to *RPS9* and calibrated to non induced iPSCs. Blue: -A-2; green: -A-4; pink: -A-5. Values are means of technical triplicates in three biological replicates . Statistical significance was determined by unpaired two-tailed Student's t-test: *p ≤ 0.01, **p ≤ 0.001, ***p ≤ 0.0001.

**(B)** Start of induction of the induced transcript 48h before differentiation. Heatmaps show bisulfite sequencing results for the analyzed amplicon on day 7 of trilineage differentiation in cell line DSMZi017-A-2. CpGs are in columns, reads are in rows, blue indicates non-methylated CpGs, red indicates methylated CpGs. Percentage of overall DNA-methylation and number of reads are given in the plots.

**(C)** Induction stopped on day 4 of trilineage differentiation. Heatmaps show bisulfite sequencing results for the analyzed amplicon on day 7 of trilineage differentiation in cell line DSMZi017-A-2. CpGs are in columns, reads are in rows, blue indicates non-methylated CpGs, red indicates methylated CpGs. Percentage of overall DNA-methylation and number of reads are given in the plots.

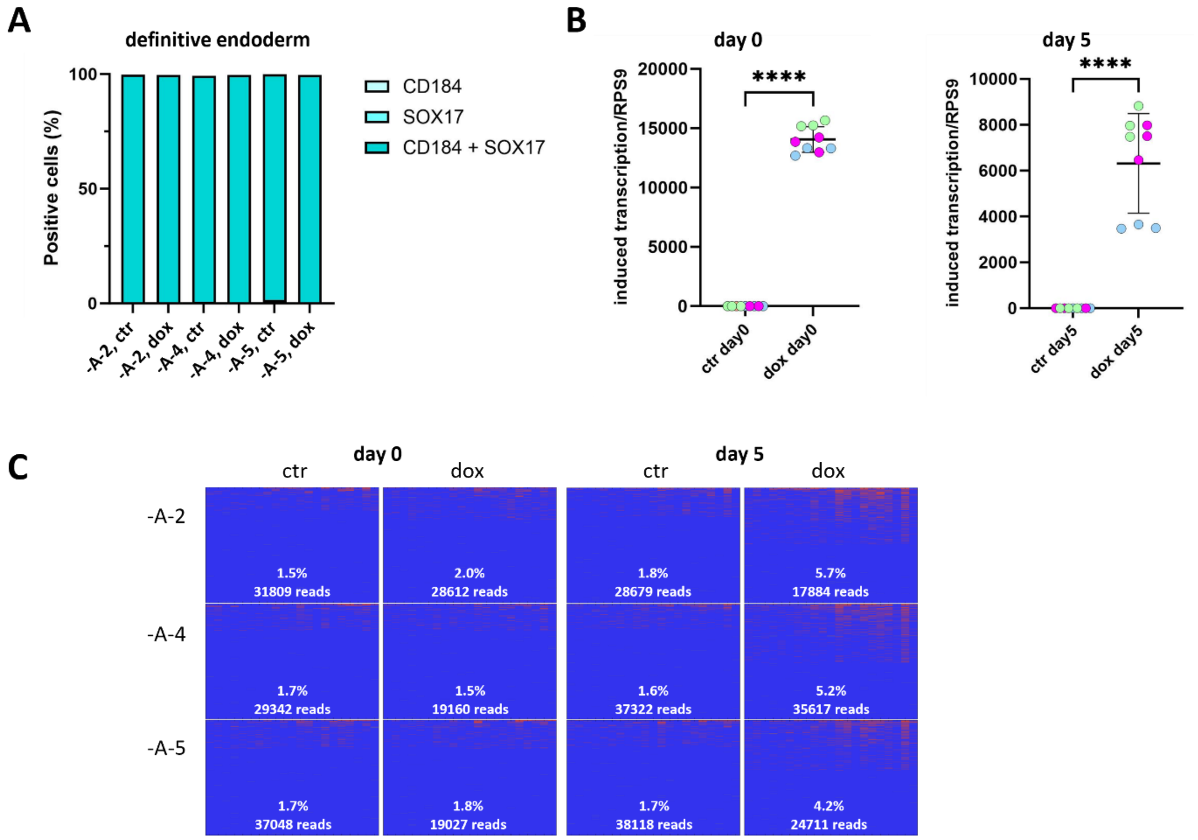

**Supplementary Figure S5. Differentiation into definitive endoderm in clonal lines DSMZi017-A-2/-A-4/-A-5.**
**(A)** Successful differentiation into definitive endoderm was confirmed on day 5 by flow cytometry for the endoderm-specific markers CD184 and SOX17.
**(B)** Induction of the transcript across the PWS-SRO was confirmed by RT-qPCR on day 0, since the iPSCs were induced by doxycycline prior differentiation, and on day 5 (end of differentiation). Induction of the inducible transcript resulted in a significantly higher level of the transcript in doxycycline-treated cells compared to the control (Student t-test, **** p≤0.0001). Expression was normalized to *RPS9* and calibrated to non-induced iPSCs. Blue: DSMZi017-A-2, green: -A-4, pink: -A-5. Values of technical triplicates are shown.
**(C)** Heatmaps showing results of bisulfite sequencing of the *SNRPN* amplicon on day 0 (left panel) and day 5 (right panel) in differentiation to definitive endoderm.

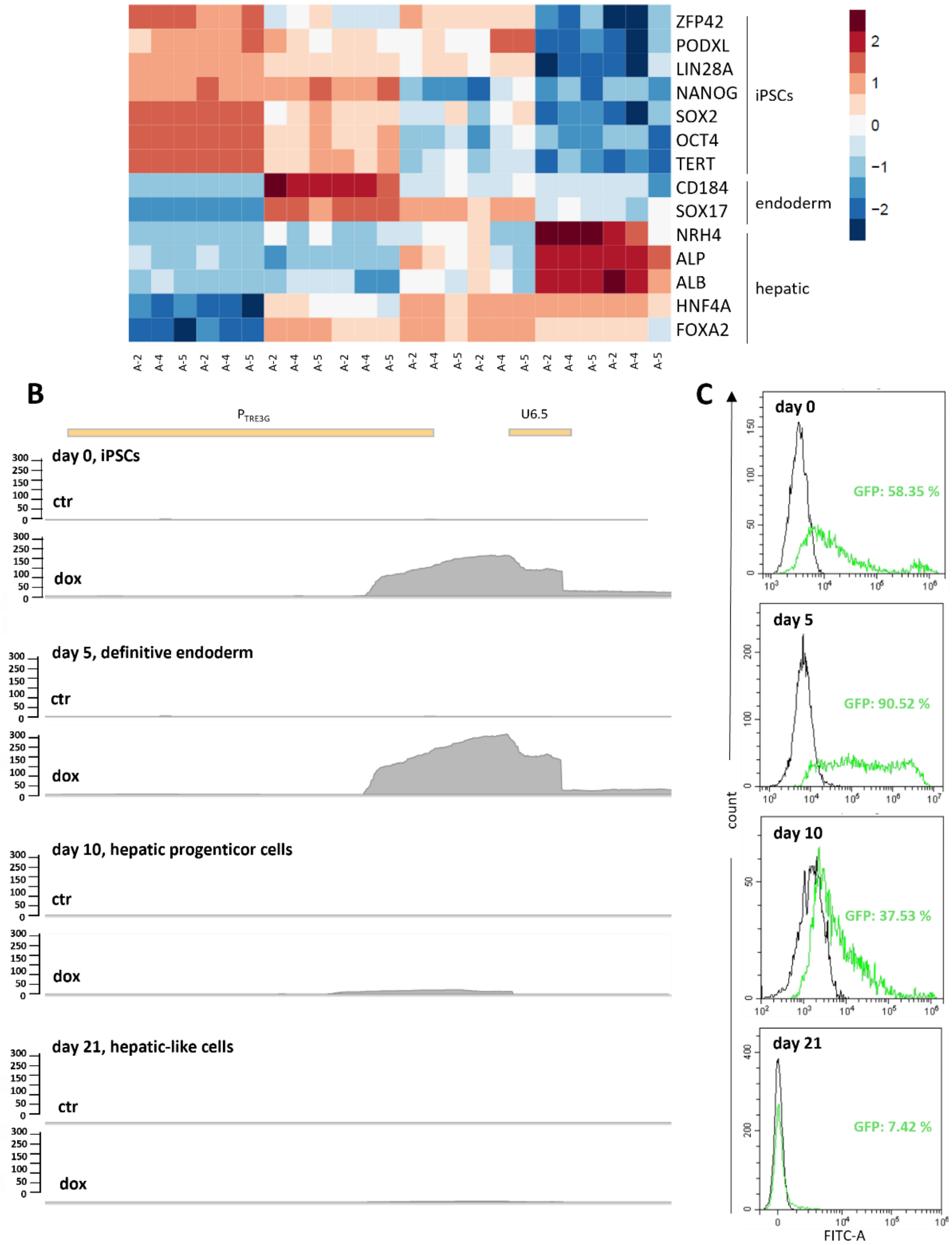

**Supplementary Figure S6. Differentiation into hepatic-like cells in clonal lines DSMZi017-A-2/-A-4/-A-5.**
**(A)** Heatmap of RNA sequencing data, representing the expression levels of specific markers for iPSCs, endodermal cells and cells from the hepatic lineage in cells untreated (ctr) or treated with doxycycline (dox). The color scale on the right represents relative expression levels, with colors ranging from low (blue) to high (red) expression.
**(B)** Genome browser-style plots display RNA sequencing read coverage for the inducible transcript region (pTRE promoter and U6.5 exon) in clonal iPSC line DSMZi017-A-2 at day 0 (iPSCs), day 5 (definitive endoderm), day 10 (hepatic progenitor cells), and day 21 (hepatic-like cells), comparing untreated and doxycycline-treated condition
**(C)** Flow cytometry data of GFP expression in clonal cell line DSMZi017-A-2 on days 0, 5, 10, and 21. Untreated control (no doxcyclin, black) defined the GFP-negative population; GFP-positive cells in doxycycline-treated samples are shown in green, with percentages indicated for each time point.

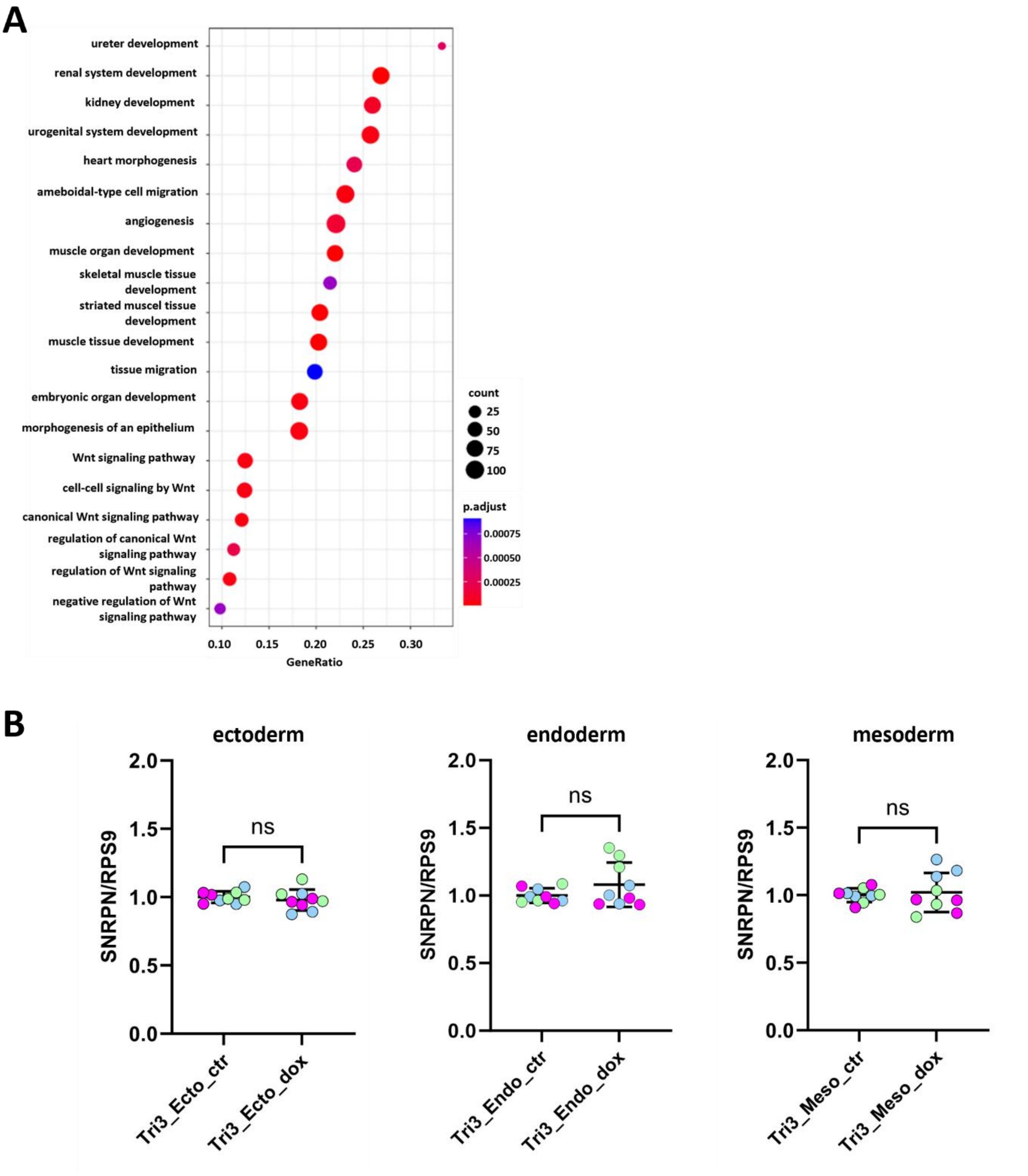

**Supplementary Fig. S7. Gene expression analysis.**

(A) Differential gene expression analysis was performed using DESeq2 on RNA sequencing data from three biological replicates (DSMZi017-A-2/-4/-45) per condition. Gene Ontology (GO) enrichment analysis was performed using the clusterProfiler R package, focusing on significantly enriched terms in the categories of biological process (BP) and molecular function (MF). Enrichment was based on differentially expressed genes (adjusted $p < 0.05$) identified by DESeq2.

(B) Results from trilineage differentiation on day 7 with stopped induction on day 4 of differentiation. DSMZi017-A-2: blue, DSMZi017-A-4: green, DSMZi017-A-5: pink. Gene expression was normalized to *RPS9* and calibrated to non-induced iPSCs. Error bars represent standard deviation. Statistical significance was determined using Student’s t-test; n.s. = not significant

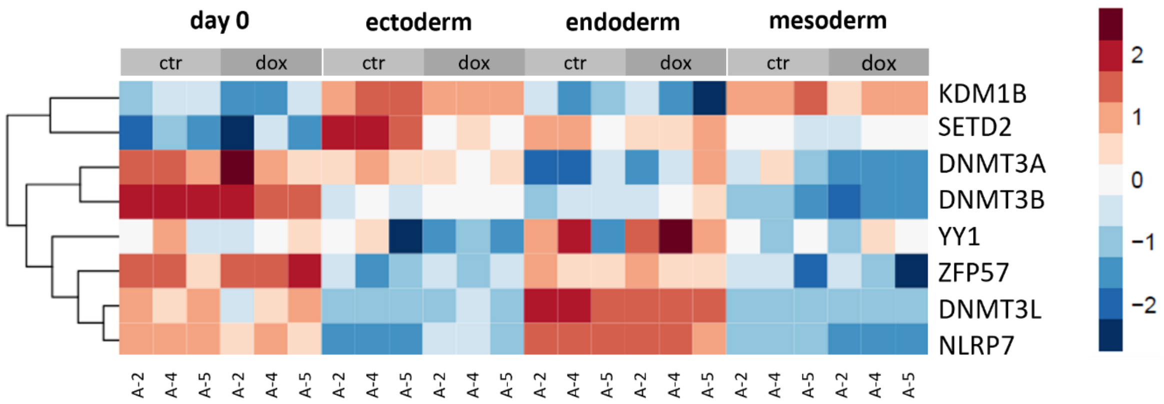

**Supplementary Figure S8. Expression of oocyte-associated factors implicated in imprint establishment during trilineage differentiation.**

Heatmap of RNA sequencing data showing expression levels of selected factors associated with de novo DNA methylation in the oocyte. Data of undifferentiated iPSC (day0) and the three germ layers (day 7) are shown for clonal cell lines DSMZi017-A-2/4/5, either untreated (ctr) or doxycycline-treated (dox). The color scale indicates relative expression levels, from low (blue) to high (red).

**Supplementary Table S1.** Antibodies

| Nuclear pluripotency markers (immunofluorescence) | | | | |
|---|---|---|---|---|
| | Dilution | Manufacturer | Catalog no | RRID |
| Nanog (D73G4) rabbit anti-human | 1:200 | Cell Signaling Technology | # 4903 | AB_10559205 |
| Oct4A (C30A3) rabbit anti-human | 1:200 | Cell Signaling Technology | # 2840 | AB_2167691 |
| Sox2 (D6D9) rabbit anti-human | 1:200 | Cell Signaling Technology | # 3579 | AB_2195767 |
| IgG Alexa Fluor 488 goat anti-rabbit | 1:1000 | Cell Signaling Technology | # 4410 | AB_1904025 |
| Surface pluripotency markers (flow cytometry) | | | | |
| | Dilution | Manufacturer | Catalog no | RRID |
| IgG1, PE | 1:25 | Miltenyi Biotec | # 130-113-438 | AB_2733893 |
| SSEA-4, PE | 1:25 | Miltenyi Biotec | # 130-122-958 | AB_2811416 |
| TRA-1-60, PE | 1:25 | Miltenyi Biotec | # 130-122-965 | AB_2801990 |
| TRA-1-81-PE | 1:25 | Miltenyi Biotec | # 130-123-334 | AB_2802035 |
| Differentiation markers (flow cytometry) | | | | |
| | Dilution | Manufacturer | Catalog no | RRID |
| CD140b, APC | 1:50 | Miltenyi Biotec | # 130-121-128 | AB_2783953 |
| CD144, FITC | 1:50 | Miltenyi Biotec | # 130-123-932 | AB_2819545 |
| CD144, PE | 1:50 | Miltenyi Biotec | # 130-118-358 | AB_2751492 |
| CD184, APC | 1:50 | Miltenyi Biotec | # 130-120-778 | AB_2752192 |
| PAX-6, APC | 1:50 | Miltenyi Biotec | # 130-123-328 | AB_2819477 |
| SOX2, FITC | 1:50 | Miltenyi Biotec | # 130-120-790 | AB_2784459 |
| SOX2, PE | 1:50 | Miltenyi Biotec | # 130-121-0530 | AB_2784460 |
| SOX17, VioB5151 | 1:50 | Miltenyi Biotec | # 130-111-147 | AB_2653496 |
| SOX17, PE | 1:50 | Miltenyi Biotec | # 130-111-032 | AB_2653493 |

**Supplementary Table S2.** Primer and guideRNA. Number in brackets at RT-PCR primers refer to numbers given in Fig. 1.

| Genotyping (after gene editing) | |
|---|---|
| Primer | Sequence (5'-3') |
| AS-SRO_LS_15F | CTGGGAGGTCTGTGAATGGAAA |
| AS-SRO_LS_15R | GGTTCACTAAACGAGCTCTGCT |
| AS-SRO_RS_15F | CTTTCCGTACCACTTCCTACCC |
| AS-SRO_RS_15R | GGTCTCAAACTCCTGACCTCAG |
| AS-SRO_region_F | AACTCAGAAGGCAACATTCCCT |
| AS-SRO_region_R | ACTGTGCTCTCTTCCCTTTCTG |
| Sanger sequencing | |
| Primer | Sequence (5'-3') |
| ASSRO_Guide_Seq_L_F | AACTCAGAAGGCAACATTCCCT |
| ASSRO_Guide_Seq_L_R | GGTCTGGGATTGAGTTTATGCC |
| ASSRO_Guide_Seq_R_F | TTCGGAGAGTTACCCAGGTGTA |
| ASSRO_Guide_Seq_R_R | ACTGTGCTCTCTTCCCTTTCTG |
| RT-PCR | |
| Primer | Sequence (5'-3') |
| (1) PTRE-F | CTTTCCGTACCACTTCCTACCC |
| (2) PTRE-R | TAACCACAGGAGGACTTGGAGA |
| (2a) U6.5_F | TTGTCTGCATTCAAGAAAGAT |
| (3) 35kb_1-F | CAGAGTAGCCCTGGAAACATGT |
| (4) 35kb_1-R | ACTGTGCTCTCTTCCCTTTCTG |
| (5) 35kb_3-F | GGCTGCAGATCCCTTCTATACC |
| (6) 35kb_3-R | TGTACTTTTCCGTGGCCTACTC |
| (7) SNRPN_Exon_1_F | CTGACGCATCTGTCTGAGGAG |
| (8) SNRPN_Exon_1_R | TTCCTCGCTACTCCAATATGGC |
| GAPDH_F | TGCACCACCAACTGCTTAGC |
| GAPDH_R | GGCATGGACTGTGGTCATGAG |
| RT-qPCR (inducible transcript and SNRPN expression | |
| Primer | Sequence (5'-3') |
| PTRE_F | CTTTCCGTACCACTTCCTACCC |
| SNRPN_Exon_1_F | CTGACGCATCTGTCTGAGGAG |
| SNRPN_Exon_3_R | TTCCTCGCTACTCCAATATGGC |
| RPS9_F | GGGAAGCGGAGCCAACATG |
| RPS9_R | GTTTGTTCCGGAGCCCATACT |
| Guide RNA (sgRNA) | |
| Primer | Sequence (5'-3') |
| sgRNA_A_left_ASSRO | TACTAATGCACACTCAAGTG |
| sgRNA_Laura_right_ASSRO | TGCATCCTCACGAATTCCCT |
